# A spreading, multi-tissue wound signal initiates whole-body regeneration

**DOI:** 10.1101/2024.12.10.627832

**Authors:** Catriona Breen, Mansi Srivastava

**Affiliations:** Department of Organismic and Evolutionary Biology, Museum of Comparative Zoology, Harvard University, Cambridge, MA 02138

**Keywords:** ERK, regeneration, wound signaling, muscle, stem cells

## Abstract

Animal bodies are equipped to sense injuries and respond, and rapid activation of ERK signaling at wound sites has emerged as a recurring mechanism across metazoans. During whole-body regeneration, distant species including cnidarians, planarians and echinoderms activate ERK at wounds to drive early transcriptional changes. However, it is unclear to what extent these are distinct phenomena, utilizing a common effector pathway but with different upstream inputs and downstream functions, or are more deeply alike, sharing upstream inputs, activating cell type(s), or spatiotemporal dynamics. To facilitate thorough comparisons across animals, we examined wound-induced ERK during whole-body regeneration in the acoel *Hofstenia miamia*. ERK signaling is induced at wound sites within 10 minutes post-amputation and peaks in activity within one hour. ERK is required for head and tail regeneration and inhibiting ERK suppresses the expression of wound-induced genes, which are themselves required for regeneration. Quantification of spatial dynamics revealed that active ERK spreads rapidly away from the wound site during the first 30 minutes post-amputation and continues to expand in space through 3-6 hours, albeit at lower levels. Multiple cell types activate ERK including neoblasts (stem cells) and muscle, and ERK is spatially dynamic within both cell types, however, ERK activation in muscle does not require neoblasts. Finally, *neuregulin-1*, a putative ligand produced exclusively by muscle cells, is required for ERK activation at wounds. Altogether, our data identify a key signaling role of muscle cells in driving ERK activation in multiple cell types after injury, and describe a spatial spreading phenomenon with parallels to dynamic ERK patterns in other systems.

## INTRODUCTION

Whole-body regeneration, the process through which some animals are able to restore entire body axes and to replace all missing cell types, is observed across distantly related animal lineages (Bely, 2010). When cut in half, animals such as cnidarians, planarians, acoels and annelids suffer disruption to many tissues including muscle, nervous system, gut, epidermis, and in some cases, stem cells. Successful regeneration involves coordinated action by cells of such diverse types located proximally to the injury site as well as cells located distally, resulting in systemic responses (Fan et al., 2023; Hernroth et al., 2010; Payzin-Dogru et al., 2024; Rodgers et al., 2017; Wenemoser and Reddien, 2010). A key early action cells must take in response to injury is to initiate changes in gene expression. For this to occur, some stimulus associated with injury must be transmitted by proteins present before the injury to shape transcriptional regulation. The extracellular signal regulated-kinase, or ERK, signaling pathway has been found to mediate this transmission: ERK signaling is activated upon injury and required for rapid transcriptional responses in distantly related animal species that display different regeneration capacities (Figure 1A) (DuBuc et al., 2014; Fan et al., 2023; Grose et al., 2002; Johnston et al., 2021; Kaloulis et al., 2004; Martin and Nobes, 1992; Owlarn et al., 2017; Tasaki et al., 2011; Tomasso et al., 2023; Tursch et al., 2022; Wolff, 2020; Zhang et al., 2024). Furthermore, ERK signaling is upstream of the wound-induced upregulation of orthologous genes across species, including both highly regenerative species such as planarians and those with limited regenerative capacity such as mice. This raises the possibility that although whole-body regeneration presents unique challenges, they are addressed beginning with ancient, conserved wound response processes. To assess this hypothesis at greater resolution, it is important to determine whether wound-induced ERK signaling is mediated and achieves its functions by similar or different means across species.

**Figure 1.**
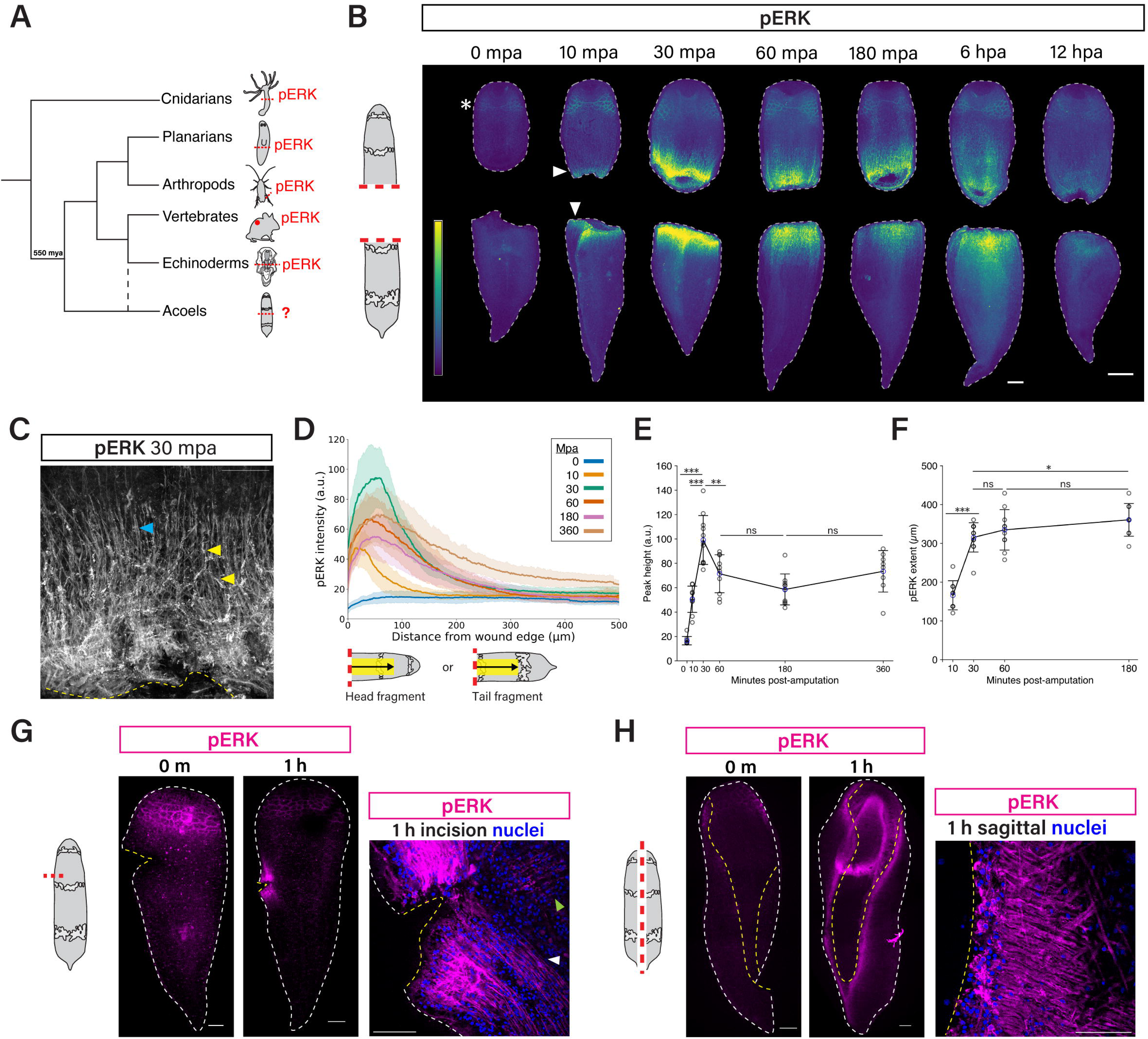
Wounding causes rapid activation, then spreading, of ERK signaling in *Hofstenia*. (A) Phylogenetic tree depicting animal taxa that include at least one species for which wounding has been shown to activate ERK signaling in the wound region within 6 hpa. Dashed line indicates suggested alternate placement of Xenacoelomorpha. *Mus, Patiria* larva and *Periplaneta* schematics modified from PhyloPic (credits: *Periplaneta* Thomas Hegna, *Patiria* T. Michael Keesey). *Hofstenia, Hydra* and planarian schematics modified from (Ramirez, 2020). (B) ERK signaling is activated at head-and tail-facing wound sites upon transverse amputation. phospho-ERK (pERK) is detected at wound sites within 10 mpa (arrowheads), is highly enriched in the wound region through 6 hpa, and is modestly enriched in this region at 12 hpa. pERK is also present in brain neurite bundles (*, 0 mpa head fragment). (C) High-magnification view of 30 mpa head fragment wound edge showing pERK^+^ muscle fibers (blue arrowhead) and neurite bundles (yellow arrowheads). (D) Quantification of pERK fluorescence intensity along the proximal-distal axis over time following transverse amputation (n >= 8 pooled head and tail fragments per time point). Shading shows standard deviation. Schematics below graph show quantified area. (E) Profile peak height, or fluorescence intensity at the location of greatest pERK intensity, at each time point. (F) Extent of pERK staining (distance at which profile slope falls below a threshold) at 10, 30, 60 and 180 mpa. Blue circles in E, F, mean; error bars, standard deviation. (G) pERK IF at small incision wounds 0 and 1 hpa. Inset shows pERK^+^ body wall muscle fibers at the injury site, both longitudinal (white arrowhead) and circumferential (green arrowhead). (H) pERK IF at a sagittal amputation wound 0 and 1 hpa. Inset shows pERK^+^ circumferential body wall muscle fibers at the wound edge. mpa, minutes post-amputation, hpa, hours post-amputation. Gray or white dotted lines show fragment outline; yellow dotted lines in (C, G, H) show wound edge. * p < 0.05, ** p < 0.01, *** p < .001, ns, not significant, two-sided Welch’s t-test. Adjusted p-values for pERK peak height: 0 vs 30 mpa 3.37e-7, 10 vs 30 mpa 1.37e-5, 30 vs 60 mpa 4.12e-3, 30 vs 180 mpa 6.51e-5, 60 vs 180 mpa .058, 180 vs 360 mpa .057. Adjusted p-values for pERK extent: 10 vs 30 mpa 1.47e-7, 30 vs 60 mpa 0.334, 60 vs 180 mpa 0.319, 30 vs 180 mpa .0347, 10 vs 180 mpa 8.27e-8. Scale bars 100 µm (B, G, H), 50 µm (C, insets G, H).

Within one hour of amputation, ERK signaling is robustly activated at the amputation site in planarians, the cnidarians *Hydra* and *Nematostella*, and sea star larvae (*Patiria*), and is required for whole-body regeneration in each (DuBuc et al., 2014; Fan et al., 2023; Johnston et al., 2021; Kaloulis et al., 2004; Owlarn et al., 2017; Tasaki et al., 2011; Tomasso et al., 2023; Tursch et al., 2022; Wolff, 2020). Injury-induced, local ERK activation is also observed in cases of more limited regeneration including appendage regeneration in an insect, tail fin regeneration in zebrafish, and skin regeneration in spiny mouse, as well as following skin injury in *Mus musculus*, which heals skin wounds but does not regenerate complete skin (Grose et al., 2002; Martin and Nobes, 1992; Tomasso et al., 2023; Yoo et al., 2012; Zhang et al., 2024). Despite the similarly rapid timing of ERK activation in these species, whether the same upstream cues trigger initial ERK activation is not fully understood. For example, reactive oxygen species (ROS) and calcium signaling promote rapid ERK activation at injuries in *Hydra;* in planarians, ROS positively regulate ERK at a later time point (1 day post-amputation), but whether ROS or calcium signaling act on ERK within the first hours post-wounding is unknown (Jaenen et al., 2021; Tursch et al., 2022). Conversely, the epidermal growth factor receptor (EGFR) ortholog *egfr3* positively regulates ERK activation in planarians, but whether receptor tyrosine kinase signaling is involved in *Hydra* is unknown. While the signals upstream of ERK are well-studied during blastema formation and growth in vertebrate skin and appendage regeneration, in these species the cues that trigger the earliest ERK activation following injury, within preexisting cells, are less well characterized (Farkas et al., 2016; Poss et al., 2000; Wen et al., 2022).

The spatial dynamics and cell type context of wound-induced ERK are also not well understood in all cases. Planarian ERK activity following amputation forms a wave, in which ERK activity is initiated close to the wound edge but occurs more and more distally over time (Fan et al., 2023). However, whether other species display this pattern has not been investigated. In terms of cell type context, in injured spiny mouse ear skin a wide range of cell types activate ERK including epidermal cells, neurons, chondrocytes and muscle cells, raising the possibility that cells of many types activate ERK in other species as well, but this is yet to be determined. These aspects of ERK as a wound signal are pertinent to the downstream biological role of ERK in regeneration, as cells make distinct contributions to regeneration depending on their cell type and location. Furthermore, the spatial, cellular and molecular dimensions of ERK activation at wounds can provide rich substrate for evolutionary comparison, building our understanding of shared, core elements of an animal body’s response to a wound.

Given the potential conservation of ERK as a wound response upstream of early transcriptional changes after wounding, we wanted to test the hypothesis that ERK serves that role in more distantly-related animal lineages. We focused on the acoel *Hofstenia miamia* (referred to hereafter as *Hofstenia*), which represents a lineage (Phylum Xenacoelomorpha) that is likely the sister to all other bilaterians, providing a large phylogenetic distance to systems where ERK has been investigated previously (Álvarez-Presas et al., 2024; Cannon et al., 2016; Ruiz-Trillo et al., 1999). Specifically, we sought to characterize ERK activation and its relationship to wound-induced gene expression, as well as to understand ERK’s spatial and cell type dynamics, including assessing the possibility of a wave or spreading pattern. *Hofstenia* has emerged as a new research organism for studies of whole-body regeneration (Srivastava, 2022). Additionally, previous work in this system characterized a wound-induced gene regulatory network led by the immediate early gene *egr*, which acts as a direct transcriptional regulator of other wound response genes (Gehrke et al., 2019; Ramirez et al., 2020). Altogether, the tools available in the system and the prior knowledge of the transcriptional wound response position *Hofstenia* to inform studies of ERK function in whole-body regeneration.

Here, we show that amputation triggers rapid activation of ERK signaling in *Hofstenia*. The first cells to activate ERK are those closest to the injury site, but over the first 3 hours post-injury, ERK is progressively activated in cells farther and farther from the injury site in a spreading pattern. Wound-induced ERK signaling is required for regeneration and promotes early transcriptional responses to wounding, including *egr* expression. Closely examining ERK’s progression upon injury, we found that ERK is activated and spatially dynamic within multiple cell types, but that muscle cells activate ERK independently of neoblasts (stem cells). Finally, we show that *neuregulin-1,* a putative ligand of epidermal growth factor family receptors (EGFRs) that is expressed solely by muscle cells, is required for ERK activation upon amputation. This demonstrates that muscle cells provide key signaling to other cells to launch their molecular responses to injury. Overall, our work reveals that an acoel employs a deeply conserved wound signal to elicit the transcriptional changes that initiate regeneration.

## RESULTS

### ERK signaling is rapidly activated at wound sites in *Hofstenia*

ERK signaling is wound-induced and drives rapid transcriptional responses at wounds in diverse animal species (Figure 1A). Therefore, we chose ERK as a candidate signaling pathway that might act upstream of wound-induced gene expression in *Hofstenia*. First, we confirmed that *Hofstenia* possesses genes encoding extracellular signal regulated-kinase (*erk*) and mitogen-activated protein kinase kinase (*mek*). We identified one putative *erk* ortholog and one putative *mek* ortholog, both of which are expressed in multiple major cell types including muscle cells, neoblasts and neurons in whole-animal single-cell RNA sequencing data (Figure S1A) (Hulett et al.). Fluorescent *in situ* hybridization (FISH) confirmed that *erk* is expressed in intact animals and does not appear to be restricted to specific body regions (Figure S1B).

To assess whether wounding activates ERK signaling in *Hofstenia*, we performed immunofluorescence (IF) for phospho-ERK (pERK), an indicator of active ERK signaling. Immediately after transverse amputation (0 minutes post-amputation, mpa), pERK labeled brain neurite bundles, but head and tail fragments did not show enriched pERK signal at the wound edge (Figure 1B). However, at 10 mpa, we observed enriched pERK in cells very close to the wound edge (Figure 1B, white arrowheads). At 30 mpa, we observed intense pERK signal in a broad band of cells beginning at the wound edge, including prominent, pERK^+^ body wall muscle fibers and neurite bundles extending away from the wound edge (Figure 1C). We continued to observe clear pERK enrichment in the wound-adjacent region at 1, 3 and 6 hours post-amputation (hpa) (Figure 1B). By 12 hpa, pERK was still enriched in this region, but appeared diminished in intensity compared to earlier time points.

We noted a rapid expansion of the pERK^+^ area between 10 and 30 mpa. To quantitatively describe this pattern, we measured pERK IF intensity along the anterior-posterior axis over time, plotting pERK signal (averaged across the mediolateral axis within the analyzed area) as a function of distance from the wound edge (Figure 1D, S1C). Plotting the average pERK profile of multiple fragments at each time point gave a visual representation of pERK’s spatial dynamics. This confirmed that while 0 mpa fragments have no pERK enrichment at the wound edge, beginning at 10 mpa the wound edge is enriched for pERK: profiles show a “peak” proximal to the wound. To complete a quantitative description of pERK dynamics, we compared specific characteristics of these pERK profiles between time points.

First, the average peak height of pERK profiles is greatest at 30 mpa (Figure 1E). Thus, of the time points we examined, the highest pERK level at any distance from the wound edge occurs at 30 mpa (adjusted p-value < .05 for all pairwise comparisons of 30 mpa to other time points, two-sample t-test). pERK peak height decreases from 30 to 60 mpa, and subsequently does not change significantly between 60 mpa and 3 hpa (adj. p-value = .058) or between 3 and 6 hpa (adj. p-value = .057). Second, in these profiles, pERK intensity is always greatest within 60 µm of the wound edge (Figure S1D). While distance from the wound edge to the pERK peak increases between 10 and 30 mpa, peak location is not significantly different between 30 mpa and any later time point. Our data suggest that from 30 mpa onward, the location of greatest pERK intensity remains largely the same and remains close to the wound edge.

Third, for at least the first 3 hpa, the extent of the pERK^+^ signal increases over time (Figure 1F). For each fragment, to determine how far away from the wound edge above-background pERK signal extends (“pERK extent”), we identified the distance at which the slope of the pERK profile falls within a threshold range of zero (Methods). By this measure, between 10 and 30 mpa, average pERK extent increases dramatically (difference of 119 µm, adj. p-value = 1.47e-7, two-sided Welch’s t-test), consistent with our impression that pERK “spreads” away from the wound edge. By 3 hpa, average pERK extent is more than 150 µm greater than at 10 mpa (difference of 162 µm, adj. p-value = 8.27e-8). We were unable to apply this analysis to 6 hpa profiles because some of these never become flat within the measured area –i.e., we did not capture a distal limit of above-background pERK. It is unknown whether in those fragments, pERK reaches background level at a distance beyond the measurement area used in this analysis, or whether pERK never reaches background level, and has continued to “spread” to encompass the entire body. Examining such 6 hpa fragments, we observed some faintly pERK^+^ muscle fibers quite far from the wound site (Figure S1E), consistent with the average 6 hpa pERK profile’s intensity being slightly greater than other time points’ at distances far from the wound edge. We note, however, that pERK’s pattern at 6 hpa is variable, with some fragments lacking these distal, pERK^+^ muscle fibers. Taken together, active ERK signaling spreads away from the wound edge for at least the first 3 hpa. Active ERK may continue to spread through 6 hpa in at least some fragments, but the level of this most distal pERK appears only slightly higher than baseline.

Finally, we asked whether ERK signaling is only activated following a major injury that prompts regeneration, or is activated following injury generally, including injuries that are resolved by wound healing without significant new tissue production. We observed robust pERK activation at small transverse incision wounds at 1 hpa, including dense, pERK^+^ longitudinal body wall muscle fibers as well as some pERK^+^ circumferential muscle fibers (Figure 1G). We also asked whether ERK activation requires injury across the anterior-posterior axis. After a sagittal amputation bisecting the animal across the mediolateral axis, pERK was strongly activated along the amputation wound at 1 hpa, with high pERK signal in circumferential muscle fibers extending away from the wound edge (Figure 1H). Thus, in *Hofstenia*, multiple types of wounds rapidly induce ERK signaling at and close to the injury site.

### Wound-induced ERK signaling is required for regeneration

Our immunofluorescence data indicate that pERK levels following amputation remain high through 6 hours, but diminish by 12 hpa. This, and our interest in functions of wound-induced ERK specifically rather than roles of ERK in later regenerative processes, led us to test the hypothesis that ERK signaling specifically within the first 3 hpa might be required to initiate regeneration. We developed a protocol to inhibit ERK phosphorylation in *Hofstenia* using the small molecule inhibitor U0126, which non-competitively inhibits MEK (Figure 2A) (Favata et al., 1998). Treating *Hofstenia* with 20 µM U0126 for 2 hours pre-amputation to allow drug entry, then for 1 hour post-amputation, inhibits ERK phosphorylation at wound sites at 1 hpa (Figure 2B).

**Figure 2.**
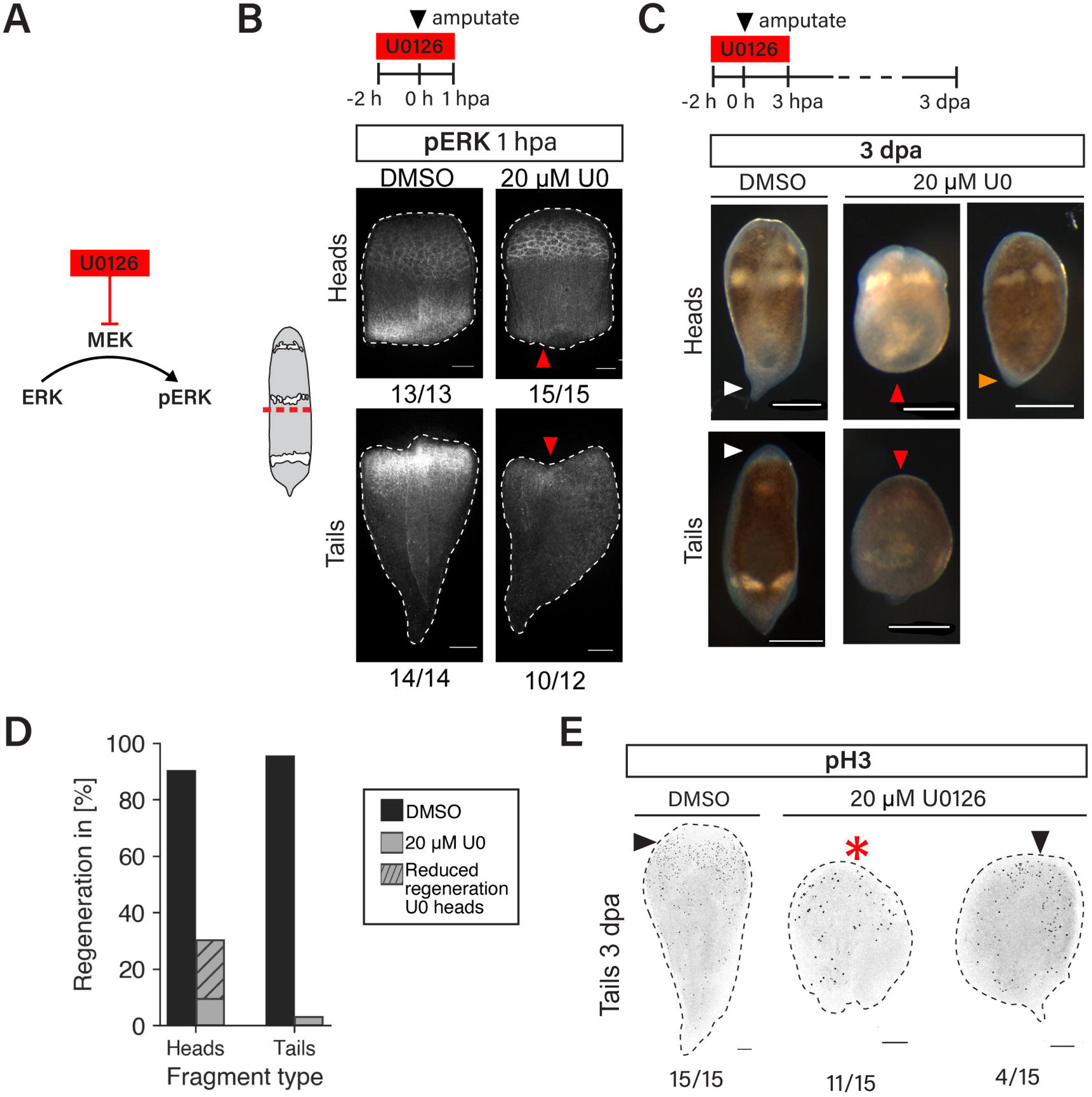
Wound-induced ERK signaling is required for regeneration. (A) Schematic of the part of the MEK-ERK pathway targeted by U0126. (B) Top, schematic showing timing of U0126 treatment. Bottom, pERK IF at 1 hpa in animals treated with DMSO (vehicle control) or U0126. pERK levels at wound sites are reduced (red arrowheads). (C) Top, schematic showing timing of U0126 treatment during regeneration assay. Animals treated with 20 µM U0126 for 2 h before and 3 h post-amputation regenerate no (red arrowheads) or reduced (orange arrowhead) visible unpigmented, new structures by 3 dpa. White arrowheads, newly formed tail and head structures of control head and tail fragments, respectively. (D) Percentages of phenotypes observed in the regeneration assay shown in (C). 29/43 U0126 heads showed no regeneration, 9/43 U0126 heads had reduced regeneration, and 33/34 U0126 tails showed no regeneration, compared to 36/40 DMSO heads and 40/42 DMSO tails with normal regeneration. Counts encompass two independent experiments, one of which is also represented in Figure S2B. (E) Tail fragments treated with 20 µM U0126 at 0 dpa do not form a normal accumulation of pH3^+^ cells in the anterior by 3 dpa (arrowheads; 4/15 U0126 tail fragments with anterior mitotic cell accumulation, compared to 15/15 control tail fragments). *, scattered pH3^+^ cells without clear anterior accumulation. Scale bars 300 µm (C), 100 µm (B, E).

Next, we asked whether temporary ERK inhibition is sufficient to inhibit regeneration. Inhibiting ERK activity using U0126 for 2 hours pre-, then 3 hours post-amputation caused stark head and tail regeneration failure (Figure 2C, D). At 3 dpa, most control head fragments (90%, n = 36/40) had a visible tail–an outgrowth with reduced pigmentation and a point at the posterior– while 29/43 head fragments (67%) treated with 20 µM U0126 failed to regenerate a visible tail, and an additional 9/43 showed a reduced degree of tail regeneration with only a small amount of unpigmented tissue. Likewise, most control tail fragments (95%, n = 40/42) showed an unpigmented anterior blastema at 3 dpa, but 33/34 tail fragments (97%) treated with 20 µM U0126 lacked a blastema. A lower U0126 dose, 15 µM, caused head and tail regeneration failure with lower penetrance (Figure S2B). To corroborate the finding that U0126-treated fragments did not regenerate new structures, we assessed cell proliferation in tail fragments, which normally show a clear accumulation of mitotic (phospho-histone H3^+^, pH3^+^) cells in the anterior at 3 dpa (Srivastava et al. 2014). Control tail fragments showed a clear accumulation of pH3^+^ cells in the anterior region at 3 dpa (15/15), but most U0-treated tail fragments showed no local accumulation of pH3^+^ cells (11/15) (Figure 2E). We conclude that ERK signaling during the first 3 hpa is required for both head and tail regeneration, including the establishment of head regeneration-associated stem cell proliferation.

To rule out the possibility that U0126 prevents regeneration due to nonspecific toxicity, we asked whether treating with U0126 at a slightly later time point also prevented regeneration. Rather than treating animals on the day of amputation (0 dpa) as above, we treated animals at 1 dpa, treating for 5 hours to match the total U0126 exposure time. Most tail fragments treated at 1 dpa with either 20 or 15 µM U0126 formed a visible new head by 3 dpa (15/20 20 µM fragments and 15/20 15 µM fragments, compared to 1/20 20 µM and 3/20 15 µM fragments treated at 0 dpa in the same experiment) (Figure S2A, B). We were able to corroborate the finding that U0126 treatment at 1 dpa does not prevent head regeneration in that at 3 dpa, most 20 µM, 1 dpa-treated tail fragments (12/15) showed a clear accumulation of pH3^+^ cells in the anterior (Figure S2C). Thus, our data indicate that exposure to 20 µM U0126 for 5 hours does not generically prevent head regeneration (Figure S2D). Furthermore, we observed no more than a 10% death rate (2/20 animals) for any group in this experiment (Table S1). Among head fragments, about half (11/20) of fragments treated 1 dpa with 20 µM U0126 failed to form a visible new tail, a similar proportion as for fragments treated 0 dpa during this experiment (14/20). This result suggests that ERK signaling may play a role in later events of tail regeneration in addition to early ones. In contrast, our data do not indicate a requirement of ERK signaling at 1 dpa for head regeneration.

To determine whether U0126-treated fragments that lacked visible new structures at day 3 had been permanently blocked from regenerating or would eventually do so, we followed a cohort of day 0, 20 µM-treated animals until 7 and 12 dpa. At both time points, many, though not all, animals regenerated a head or tail as appropriate (Figure S2E, Table S1). Therefore, ERK signaling during the first three hpa is required for regeneration at the typical rate.

### ERK drives expression of *egr*, a key regulator of the transcriptional wound response, and *egr* target genes

Previous studies of early changes in gene expression as *Hofstenia* regenerates have highlighted the importance of the transcription factor *egr*. *egr* is expressed at wound sites by 1 hpa and is required for expression of a number of other of wound-induced genes at 6 hpa as a direct transcriptional regulator (Gehrke et al). Given our findings of robust ERK activation at wound sites prior to 1 hpa and of an early requirement for ERK signaling for regeneration, we hypothesized that ERK signaling drives *egr* expression. Such a relationship would also be consistent with Egr being an immediate-early gene transcribed downstream of ERK signaling in mammalian cells in response to various stimuli, and wound-induced *egr2* expression requiring ERK signaling in planarians (Fowler et al, Sheng and Greenberg, Owlarn et al).

To test this, we first combined IF with FISH to assess whether pERK is present in cells that express *egr* at 1 hpa. We found that there is broad overlap of pERK and *egr* signal close to the wound sites of head and tail fragments observed as pERK^+^, *egr*^+^ cells (Figure 3A). Next, we inhibited wound-induced ERK signaling using U0126 and asked whether *egr* expression was affected. During experiments testing U0126 efficacy, we found that treating animals with 20 µM U0126 for 6 hours pre-amputation, then 1 hour post-amputation inhibits virtually all ERK activation at 1 hour post-amputation, in contrast to 2 hour pre-treatment which causes a clear decrease but not complete abrogation of pERK at the wound edge (Figure 2B, S3A). We therefore pre-treated animals for 6 hours with 20 µM U0126 to fully inhibit wound-induced ERK activation and assessed *egr* expression at 3 hpa (Figure 3B). *egr* expression at the wound site was diminished as compared to controls in both head and tail fragments. Therefore, we conclude that ERK promotes *egr* upregulation at the wound site upon amputation.

**Figure 3.**
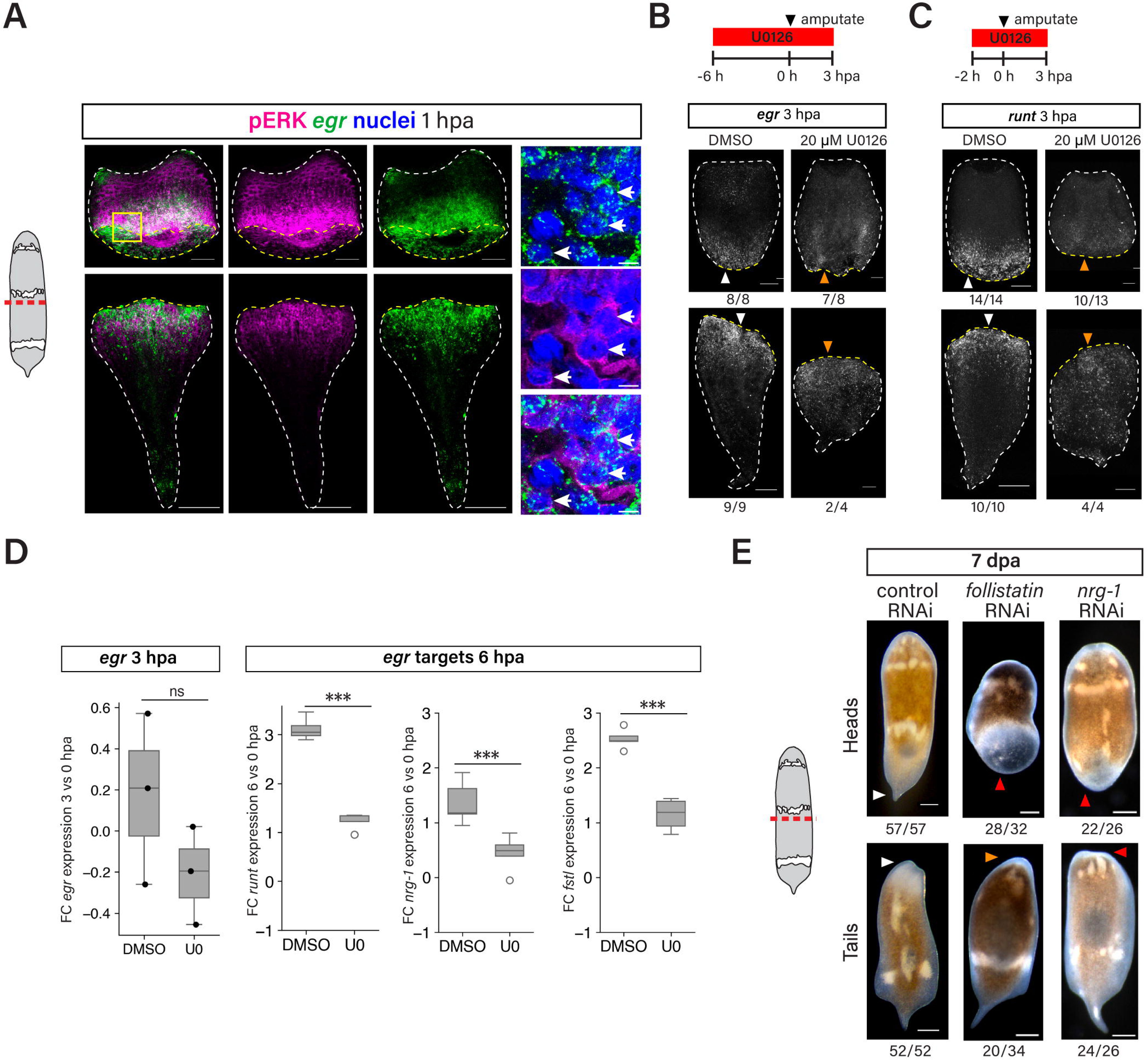
Wound-induced ERK signaling drives early transcriptional wound responses. (A) At 1 hpa, pERK and *egr* colocalize and show broad overlap of domain in head and tail fragments. Box depicts area represented in high-magnification images. Arrowheads, pERK^+^ *egr*^+^ cells. Whole-fragment and high-magnification images are single optical slices. (B, C) Top, schematics showing timing of U0126 treatment. Bottom, U0126-treated animals show diminished wound-induced *egr* (B) and *runt* (C) expression at 3 hpa. Arrowheads, wound site expression of respective genes; orange arrowheads, reduced wound site expression compared to controls. White dotted lines in (A-C) show fragment outline; yellow dotted lines show wound edge. (D) qPCR measurements of *egr* upregulation at 3 hpa and *runt*, *nrg-1* and *follistatin* upregulation at 6 hpa in control and U0126-treated fragments. Graphs show difference between log2 fold expression at 3 hpa and log2 fold expression at 0 hpa (or 6 hpa vs 0 hpa for *runt*, *nrg-1*, *follistatin*). *, p < 0.01 (two-sided Welch’s t-test). (E) *follistatin* and *nrg-1* are required for regeneration. White arrowheads, normal regeneration (57/57 control head fragments, 52/52 control tail fragments); orange arrowhead, reduced head regeneration (20/34 *follistatin* RNAi tail fragments); red arrowhead, no regeneration (28/32 *follistatin* RNAi head fragments, 24/26 *nrg-1* RNAi head fragments, 22/26 *nrg-1* RNAi tail fragments). * p < 0.05, two-sided Welch’s t-test. P-values: 0.552, 6.91e-7, 4.00e-3, 4.49e-5 for *egr, runt, nrg-1* and *fstl,* respectively. Scale bars 100 µm (A-C), 5 µm (insets in A), 200 µm (E).

If ERK signaling promotes wound-induced expression of *egr*, which itself drives expression of several additional wound-induced genes, we might expect that inhibiting ERK activity would prevent expression of those *egr* targets. *runt* is a transcription factor expressed downstream of *egr,* and among the validated *egr* targets, it is upregulated the earliest, beginning at 3 hpa (Gehrke et al., 2019; Ramirez et al., 2020). Indeed, treating animals with U0126 caused reduced *runt* expression at the wound site at 3 hpa (Figure 3C). To corroborate these FISH data, we performed quantitative PCR (qPCR) for *egr*, *runt* and two additional *egr* targets which are upregulated at 6 hpa, *follistatin* (*fstl*) and *neuregulin-1* (*nrg-1*) (Gehrke et al). Using the drug treatment schemes described above (6 hour pre-treat, 3 hour post-treat for *egr*; 2 hour pre-treat, 3 hour post-treat for *runt, nrg-1* and *fstl*), we isolated wound sites at 0 hpa and 3 (*egr*) or 6 hpa (*runt, nrg-1, fstl*). For each gene, we determined the difference in expression between 0 and 3 or 6 hpa in each condition (control or U0126-treated). For *egr*, though not statistically significant, the mean magnitude of upregulation from 0 to 6 hpa was lower in U0126-treated fragments than DMSO-treated (Figure 3D). For *runt, nrg-1* and *fstl*, upregulation from 0 to 6 hpa was significantly lower in U0126-treated fragments. This demonstrates that wound-induced ERK signaling promotes expression of *egr* as well as *egr* target genes that are also expressed within the first several hpa.

*egr* and *runt* are both known to be required for regeneration in *Hofstenia*; *runt* was also shown to positively regulate upregulation of *nrg-1* upon amputation, suggesting that *runt*, like *egr*, may promote wound-induced gene expression changes broadly (Gehrke et al). However, the roles of *nrg-1* and *fstl* in regeneration have not previously been functionally tested in *Hofstenia*. Therefore, we performed RNA interference (RNAi) to deplete either *fstl* or *nrg-1* mRNA, amputated animals and assessed regeneration outcomes. Both *fstl* and *nrg-1* RNAi head fragments failed to regenerate tails (*fstl* 88%, n = 28/32, *nrg-1* 85%, n = 22/26) (Figure 3E). *fstl* RNAi tail fragments regenerated head blastemas with reduced size (59%, n = 20/34), and *nrg-1* tail fragments showed complete head regeneration failure (92%, n = 24/26). Alongside past work on *egr* and *runt*, these results highlight that multiple ERK-dependent, wound-induced gene expression events are required for regeneration.

### ERK is activated in muscle and neoblasts, but ERK activation in muscle does not require neoblasts

Having identified downstream effects of wound-induced ERK signaling in *Hofstenia*, we were interested in further characterizing the process of ERK activation itself—in particular, pERK’s dramatic expansion in space between 10 and 30-60 mpa (Figure 1F). We first sought to understand how cell type might shape this phenomenon.

In *Hofstenia*, several major cell types including muscle cells, neoblasts, neurons, and epidermal cells show expression of *egr* within 1 hpa (Gehrke et al). We asked whether wounding activates ERK signaling within these cell types by combining IF for pERK with FISH for markers of neoblasts (*piwi-1*) or muscle cells (*tropomyosin*). At 0 mpa, we did not detect pERK within *piwi-1^+^* or *tropomyosin^+^* cells at the wound site, but at 10 mpa as well as 1 hpa, we detected pERK in *tropomyosin^+^* as well as *piwi-1^+^*cells at head and tail fragment wound sites (Figure 4A). We conclude that upon wounding, ERK signaling is rapidly activated in at least both muscle cells and neoblasts.

**Figure 4.**
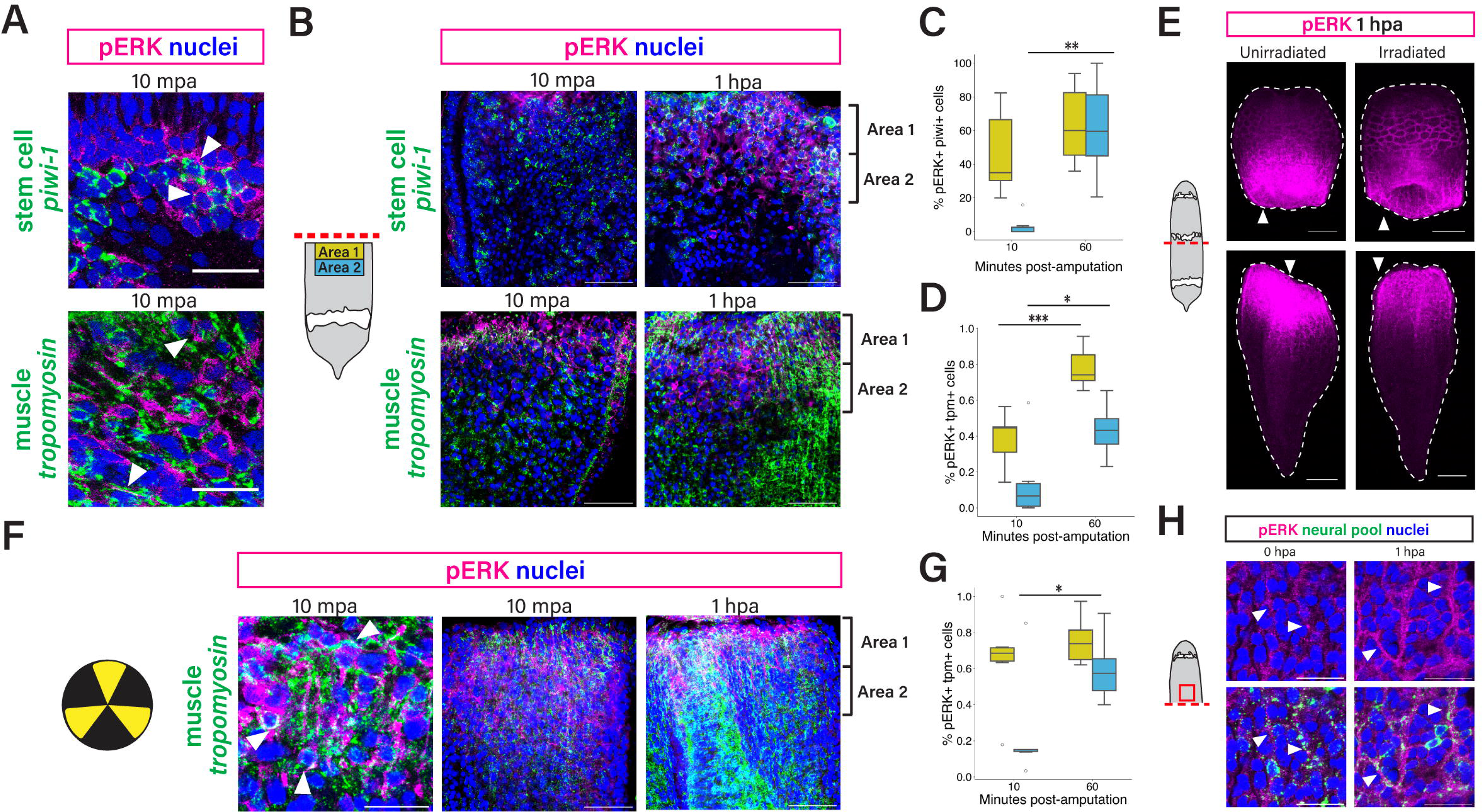
ERK is activated in muscle and neoblasts, but ERK activation in muscle does not require neoblasts. (A) ERK signaling is activated in muscle cells and neoblasts by 10 mpa (arrowheads) in both head and tail fragments. Images shown are of head fragments. (B) Left, schematic showing counting areas used for colocalization analysis: area 1 spans 0 – 50 µm away from the wound edge and area 2 spans 50 – 100 µm away. Right, representative images used for analysis of pERK cell type localization at 10 and 60 mpa. (C) Fraction of *piwi-1*^+^ cells (neoblasts) in each area that are pERK^+^ at 10 and 60 mpa. In (C), (D) and (G), yellow boxes represent cells in area 1 and blue boxes represent cells in area 2. (D) Fraction of *tropomyosin*^+^ (muscle) cells in each area that are pERK^+^. P-values: 6.35e-4 (area 1), .028 (area 2), two-sided Welch’s t-test. (E) Neoblasts are not required for overall ERK activation at 1 hpa (n = 5/5 unirradiated heads, 2/2 unirradiated tails, 8/8 irradiated heads and 6/6 irradiated tails). Images are maximum intensity projections through entire fragment. (F) Left image, ERK is activated in muscle at 10 mpa in irradiated fragments (arrowheads). Center and right images, representative images used for analysis of pERK, *tropomyosin* colocalization in irradiated animals at 10 and 60 mpa. (G) Fraction of *tropomyosin*^+^ (muscle) cells in each area that are pERK^+^ in irradiated animals. (H) FISH for neural marker pool combined with pERK IF. pERK signal within neurons appears to increase in intensity between 0 and 1 hpa, suggesting wounding may activate ERK in neurons. Images shown are of wound-adjacent region of a head fragment. * p < 0.05, ** p < 0.01, *** p < 0.001, two-sided Welch’s t-test. Error bars show standard deviation. Scale bars, 20 µm (A, left image in F, H), 50 µm (B, center and right images in F), 100 µm (E).

Next, we sought to determine whether the spatial spreading of active ERK that we observed at organismal scale (Figure 1) comprises spatially dynamic ERK activity within multiple cell types—in other words, whether ERK activity in one or multiple cell types gives rise to this overall pattern. To do this, we analyzed colocalization of pERK and *piwi-1* or *tropomyosin* within two regions of fragments: “area 1” beginning at the wound edge and extending 50 µm towards the anterior (for heads) or posterior (for tails), and “area 2” spanning 50 to 100 µm away from the wound edge (Figure 4B). We scored *piwi-1^+^* and *tropomyosin^+^*cells as either pERK^+^ or pERK*^-^*, and while binary scoring undoubtedly masks real variation among cells’ pERK levels, this approach captured population-level differences in ERK activation between wound site-proximal and more distal cells. Considering neoblasts, between 10 and 60 mpa, the fraction of *piwi-1^+^*cells in area 2 that are pERK^+^ increases significantly, from near 0 to about 60% (p-value = 3.92e-3, two-sided Welch’s t-test) (Figure 4C). In other words, at 10 mpa, neoblasts primarily within 50 µm of the wound site have active ERK signaling, while by 1 hpa neoblasts 50 -100 µm from the wound site have also activated ERK. We describe this as active ERK spreading away from the wound site over time among neoblasts. Considering muscle, the fraction of pERK^+^, *tropomyosin^+^* cells increases significantly between 10 and 60 mpa in both areas (Figure 4D). Thus, ERK activity is also dynamic within muscle cells.

To examine possible causal relationships between ERK activation in different cell types, we next assessed whether neoblasts are necessary for wound-induced ERK activation. We lethally irradiated animals and waited 7 days, which depletes all neoblasts (Srivastava et al., 2014), then amputated animals and performed IF for pERK. Irradiated head and tail fragments showed clear pERK staining at the wound site at 1 hpa, demonstrating that neoblasts are not required for overall ERK activation (Figure 4E). We repeated our *tropomyosin,* pERK colocalization analysis on irradiated fragments and observed pERK^+^, *tropomyosin^+^* cells at both 10 mpa and 1 hpa, demonstrating that muscle cells can activate ERK in response to injury in the absence of neoblasts (Figure 4F). In fact, for each area at each time point, the fraction of *tropomyosin^+^*, pERK^+^ cells was not significantly different between unirradiated and irradiated animals (zone 1 10 mpa adjusted p-value = 0.233, zone 1 60 mpa adj. p-value = 0.730, zone 2 10 mpa adj. p-value = 0.700, zone 2 60 mpa adj. p-value = 0.233, two-sided Welch’s t-test) (Figure 4D, G). This suggests that neoblasts do not play a significant role in non-cell-autonomously mediating ERK activation within muscle cells.

Finally, we also assessed whether ERK is activated in neurons upon amputation by combining pERK IF with FISH for a pool of three neural markers, *PC2, TrpC-1* and *gad-1*. In head and tail fragments at both 0 and 1 hpa, neural cell bodies and cell projections throughout the body show some pERK signal (Figure 4H, S4A). At 1 hpa, we observed that neural projections close to the wound site appear to have higher pERK signal than neural projections distant from the wound site. However, the difference in pERK intensity marking wound-proximal versus distal neural cells was subtle enough that we did not feel confident calling pERK-positive versus negative neurons. Therefore, we conclude that ERK signaling is active in neurons at 1 hpa, but cannot determine the extent to which this activation is wound-induced versus maintained from homeostatic levels.

### *Neuregulin-1,* a putative ligand produced solely by muscle cells, is required for ERK activation

Diverse upstream activators can feed into ERK signaling. In considering which proteins might bridge injury to ERK activation, we noted that the ERK spread we observe upon amputation in *Hofstenia* is reminiscent of ERK signaling waves described in other systems (Hiratsuka et al, De Simone et al, Ogura et al, Gagliardi et al). Though these phenomena are not underpinned by identical genetic or protein circuits, many are united by FGF or EGF receptor tyrosine kinase signaling activating ERK. Therefore, we hypothesized that receptor tyrosine kinase signaling might activate ERK upon amputation in *Hofstenia*. We treated animals with the small molecule SU5402, commonly used to inhibit FGF receptor activity (Mohammadi et al., 1997), and examined ERK activation at 1 hpa. At 5 µM SU5402, pERK signal at the wound edge was reduced compared to controls, and 35 µM, pERK was sharply reduced, though faintly pERK^+^ muscle fibers were visible close to the wound edge (Figure 5A). At 100 µM SU5402, wound-induced pERK was completely abolished. We measured pERK profiles of these fragments and confirmed that pERK peak height and extent decrease with increasing doses of SU5402 (Figure 5B-D). We also noted that ERK activation at 10 mpa was strongly inhibited by as little as 5 µM SU5402 (Figure S5A-D). Together, these data suggest that both initial and continued ERK activation following amputation require receptor tyrosine kinase signaling. Furthermore, SU5402 treatment strongly inhibited *egr* and *runt* expression at 3 hpa (Figure 5E, S5E). This effect appeared even stronger than that we had observed for U0126. This further supports the conclusion that ERK signaling is upstream of *egr* and *runt* expression.

**Figure 5.**
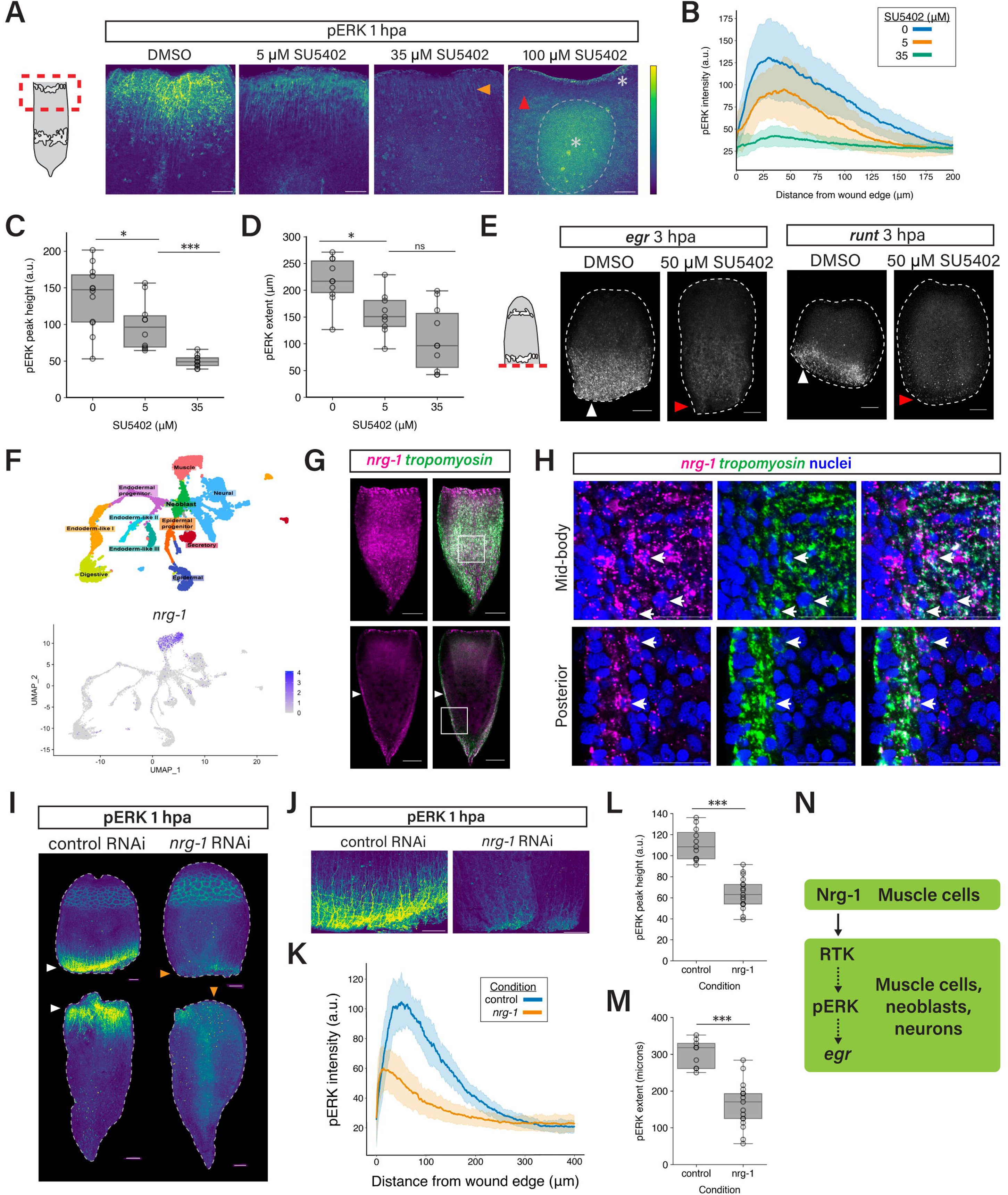
*Neuregulin-1,* a putative EGFR family ligand produced by muscle cells, is required for ERK activation. (A) pERK staining at 1 hpa in representative tail fragments treated with DMSO or increasing concentrations of SU5402. Images show maximum intensity projections of a fixed number of optical slices. Orange arrowhead, muscle fibers with low but above-background pERK signal. Red arrowhead, absence of pERK^+^ muscle fibers. In 100 µM image, * and dotted line indicate nonspecific signal. (B) Average pERK fluorescence intensity profiles of 1 hpa fragments treated with 0 (DMSO), 5 or 35 µM SU5402 (pooled heads and tails, n > 10 fragments per condition). (C, D) pERK profile peak height (C) and pERK extent (D) at 1 hpa for each SU5402 condition. Measured pERK extent in control fragments is less than measured pERK extent of untreated 1 hpa fragments (Figure 1B, F) because images for the SU5402 experiment were acquired at higher magnification and did not all encompass pERK signal beyond 200 µm of the wound edge. Therefore, for this experiment a lower slope threshold was used to call extent, yielding a relative measure of extent comparable between groups within this experiment. Adjusted p-values for pERK peak height comparisons, two-sided Welch’s t-test: .0357 for 0 vs 5 µM, 3.10e-for 5 vs 35 µM. Adjusted p-values for pERK extent comparisons: .0104 for 0 vs 5 µM , .068 for vs 35 µM . (E) *egr* and *runt* expression at 3 hpa in fragments treated with DMSO or 50 µM SU5402. 8/8 SU5402-treated heads showed sharply reduced *egr* expression as compared to controls (red arrowheads; normal *egr* expression in 5/5 control heads, white arrowheads). 4/4 SU5402-treated heads showed reduced *runt* expression as compared to controls (red arrowheads; normal *runt* expression in 10/10 control heads, white arrowheads). (F) Top, projection of intact animal scRNA-seq data from (Hulett et al., 2023) annotated with validated cluster identities. Bottom, *nrg-1* expression in the same projection of cells showing that expression is primarily within muscle clusters. (G) Double FISH for *nrg-1* and *tropomyosin* in intact animals. Top row, single optical slice in a plane encompassing body wall and peripheral muscle. *nrg-1* is expressed widely in the planes of peripheral and body wall muscle. Bottom row, single optical slice in an internal plane of the body, with peripheral and body wall muscle visible at the periphery of the body. *nrg-1* and *tropomyosin* signal overlap (white) in body wall muscle (arrowheads). Boxes show areas represented in (H). (H) *nrg-1* and *tropomyosin* mRNA colocalize (arrowheads). Top, body wall muscle cells in same view as (G) top row; bottom, posterior body wall muscle cells in same view as (G) bottom row. (I) ERK activation at 1 hpa is severely diminished in *nrg-1* RNAi animals (orange arrowheads) compared to controls (white arrowheads). (J) Wound region insets of fragments in (I). (K) Average pERK intensity profiles for control and *nrg-1* RNAi animals (pooled head and tail fragments, n = 11 control fragments and 17 *nrg-1* RNAi fragments). (L, M) pERK profile peak height (L) and pERK extent (M) at 1 hpa for control and *nrg-1* RNAi animals. Peak height and pERK extent both differ significantly between groups (adj. p-values 1.84e-8 and 2.92e-6, respectively, two-sided Welch’s t-test). (N) Model for upregulation of *egr* and *egr* target genes upon injury. Nrg-1 produced by muscle cells binds to a receptor tyrosine kinase expressed by multiple cell types, leading to ERK activation within those cell types. pERK and/or its target kinases then phosphorylate transcription factors which directly regulate expression of *egr.* Shading in (B, K) and error bars in (C, D, L, M) show standard deviation. * p < 0.05, *** p < .001, ns, not significant, two-sided Welch’s t-test. Scale bars, 50 μm (A), 100 μm (E, G, I, J), 20 μm (H).

We reasoned that a receptor-ligand pair driving ERK activation would be expressed in the intact animal so that wound-induced ERK activation could take place rapidly, without requiring new gene expression. We also reasoned that a receptor should be expressed in at least the cell types with active ERK signaling post-injury: muscle, neoblasts and neurons. Motivated by our SU5402 results and the prevalence of FGF and EGF signaling in ERK waves, we examined the expression patterns of FGF and EGF family receptors and ligands in intact *Hofstenia* in existing single-cell RNA-seq data (Hulett et al., 2023). Neuregulins are EGF family ligands that bind to ERBB receptor tyrosine kinases; *Hofstenia neuregulin-1 (nrg-1*) and a pair of putative ERBB receptor homologs, *erbb4* and *erbb4-2,* satisfied the criteria listed above (Figure 5F, S5F-H). Furthermore, *nrg-1* RNAi animals failed to regenerate (Figure 3E), although this could result from either a requirement for homeostatic *nrg-1* expression or a requirement for wound-induced *nrg-1* expression, as *nrg-1* is upregulated at wound sites at 6 hpa (Gehrke et al., 2019). *Hofstenia* possesses additional EGF family ligands, *nrg-2* and *egf*; in intact animals, *nrg-2* is expressed in the epidermal lineage and *egf* is expressed in several cell populations including muscle, neurons, endodermal progenitors and digestive cells (Figure S5H). The *Hofstenia* genome also encodes a putative FGF receptor, and it is expressed in muscle and endodermal progenitor cells of intact animals (Figure S5H). As *fgfr* is not expressed in neoblasts or neurons in single-cell RNA-seq data, we reasoned that *fgfr* could not drive ERK activation in those cell types cell-autonomously, so we did not pursue *fgfr* as a candidate. We were not able to identify any FGF ligand orthologs in the *Hofstenia* genome.

Single-cell RNA-seq data indicate that in intact animals, only muscle cells express *nrg-1,* while *erbb4* and *erbb4-2* exhibit broad expression including in muscle, neoblasts and neurons (Figure 5F, S5F-H). Using double FISH for *nrg-1* and *tropomyosin*, we confirmed that in intact animals *nrg-1* is expressed body-wide in the planes of both peripheral and body wall muscle, with little signal in the interior of the body in other planes, and that *nrg-1* and *tropomyosin* transcripts colocalize (Figure 5G, H). During regeneration, in single-cell RNA-seq data, muscle clusters remain the only clusters with *nrg-1* expression (Figure S5G). We note that while *nrg-1* is one of the wound-induced genes whose upregulation at 6 hpa we found to be downstream of ERK signaling (Figure 3D) (Gehrke et al., 2019), we hypothesized that Nrg-1 protein present in intact animals might act upstream of ERK activation.

With the goal of depleting Nrg-1 protein prior to amputation, we performed a homeostatic RNAi experiment in which we subjected animals to control or *nrg-1* RNAi for 12 days. We then amputated animals and performed IF for pERK at 1 hpa. Both control and *nrg-1* head and tail fragments showed detectable pERK signal at the wound site (14/17 *unc* and 21/22 *nrg-1*); however, the intensity and spatial extent of pERK staining appeared dramatically weakened in *nrg-1* fragments (Figure 5I, J). We quantified profiles of pERK signal and confirmed these observations (Figure 5K-M). Furthermore, to rule out the possibility that *nrg-1* RNAi animals failed to activate ERK at normal levels because *nrg-1* is required for homeostatic maintenance of muscle, which itself activates ERK, we assessed *nrg-1* RNAi animals for gross muscle loss or defects using IF for Tropomyosin. *nrg-1* RNAi animals showed no gross defects in peripheral or body wall musculature (Figure S5I). Given that neuregulins bind to EGFR family receptors in other systems (Lei et al., 2016), and that *erbb4* and *erbb4-2* are the only EGFR family receptors we identified in *Hofstenia,* we speculate that *nrg-1* activates ERK signaling through *erbb4* and/or *erbb4-2*, and that SU5402 is able to inhibit the activity of these receptors in *Hofstenia.* We conclude that a factor expressed by muscle is essential for ERK activation, which takes place not only in muscle but in multiple cell types (Figure 5N) (Hulett et al., 2023).

## DISCUSSION

Our results demonstrate that whole-body regeneration in an acoel utilizes ERK signaling, a pathway activated at injury sites in diverse animal species, to launch early transcriptional events of regeneration. If ERK activation is inhibited during the first three hours post-amputation, *Hofstenia* do not regenerate, consistent with ERK playing an important role in early steps during regeneration. We further show that while ERK is activated in multiple cell types, muscle cells play a central role in ERK activation overall: muscle cells express *nrg-1*, without which ERK activation is severely diminished. This work fills an important gap in knowledge of the mechanisms linking injury to regenerative responses in *Hofstenia*, and enables comparison of specific characteristics of wound-induced ERK signaling across species.

Our finding that ERK signaling promotes expression of *egr, runt, nrg-1* and *follistatin* upon amputation mirrors regulatory relationships observed in the wound responses of other species. An *egr* homolog is also expressed downstream of ERK following amputation in planarians and skin injury in spiny mouse (Owlarn et al., 2017; Tomasso et al., 2023; Wenemoser et al., 2012). *egr* furthermore has long been known as an immediate early gene (IEG) transcribed downstream of ERK in mammalian cells stimulated by growth factors (Fowler et al., 2011). Other mammalian ERK-dependent, immediately early genes, such as *fos* and *jun*, are also upregulated post-injury in highly regenerative species including *Hydra,* ctenophores and planarians, and it will be important to assess in the future whether these genes are also part of the ERK-dependent wound response in *Hofstenia* (Cazet et al., 2021; Mitchell et al., 2024; Naziri, 2024; Wenemoser et al., 2012). Finally, in addition to classic immediate early genes, two other genes show similarity: 1) a *runt* homolog is expressed downstream of wound-induced ERK in *Hofstenia*, planarians and sea anemones, and 2) a *follistatin* homolog is ERK-regulated in both planarians and *Hofstenia* (Johnston et al., 2021; Owlarn et al., 2017). In mammalian cells, to regulate transcription, ERK signaling alters the activity of specific transcription factors including CREB, ATF and ELK family members as well as transcription factors encoded by immediate early genes themselves (EGR, FOS and JUN) (Lavoie et al., 2020). Although only a partially overlapping list of genes is found to be downstream of ERK across species thus far, we think that *in toto* the data indicate that we can expect to find strong correspondence of mechanism driving transcriptional events downstream of ERK. This can be validated thorough comparative studies in species such as planarians, cnidarians, and acoels.

Among wound-induced genes we show to be expressed downstream of ERK in *Hofstenia,* two (*egr* and *runt*) were previously reported to cause regeneration failure when inhibited via RNAi, and we show that *follistatin* and *nrg-1* are required for regeneration as well (Gehrke et al., 2019). Thus, we propose that *Hofstenia* treated with a MEK inhibitor, U0126, for a short period surrounding the time of amputation fail to regenerate by 3 dpa because of a defective transcriptional wound response. Interestingly, by 7 dpa, many U0126-treated fragments do regenerate—however, the stark difference in extent of new tissue formation between control and U0126-treated fragments at 3 dpa indicates that U0126-treated animals are delayed by much longer than the duration of drug treatment (5 hours). This suggests that following an injury, there is a critical period during which ERK signaling must occur in order for regeneration to proceed at its typical pace, a phenomenon also reported in planarians (Owlarn et al., 2017). Why this is the case remains unknown, but it is possible that a transient, injury-associated cue is needed for ERK activation, and that if enough time passes for this cue to disappear, ERK is not activated. Nonetheless, one possible reason why temporarily ERK-inhibited *Hofstenia* eventually do regenerate is that homeostatic tissue turnover is sufficient to rebuild lost structures in a matter of days; an alternative is that regeneration-specific processes take place, but more slowly than usual.

ERK signaling that occurs as a wave or spread and is downstream of an FGF or EGF family ligand is observed both in amputated *Hofstenia* and in a number of injury or developmental contexts in other species (Aikin et al., 2020; Aoki et al., 2017, 2013; De Simone et al., 2021; Ender et al., 2022; Fan et al., 2023; Gagliardi et al., 2021; Hiratsuka et al., 2015; Ishii et al., 2021; Ogura et al., 2018). Notably, in whole-body regeneration contexts, *Hofstenia* and planarians each show spatially dynamic ERK activity within the first several hours post-injury, with ERK activation beginning closest to the wound edge (Fan et al., 2023). In planarians, at later time points, relative ERK activity (i.e. the ratio of phospho-ERK to total ERK) is greatest at locations distant from the wound edge. In *Hofstenia,* the location of greatest pERK level does not significantly change after 30 mpa and remains close to the wound edge (Figure S1D). It is difficult to directly compare the two species’ patterns because we measured IF signal using a phospho-specific ERK antibody, so our measurements correspond to absolute levels of active ERK signaling rather than relative levels.

In terms of cell type dynamics, planarian work and this study both implicate muscle cells in the process of ERK activation. Planarian longitudinal muscle fibers are required for ERK-dependent transcriptional responses in wound edge-distal cells (Fan et al., 2023), and in *Hofstenia* muscle cells are the source of a factor required for ERK activation, *nrg-1*. Given that planarian *egfr3* positively regulates ERK activation at 1 dpa (Jaenen et al., 2021), it will be interesting to know whether neuregulins or EGFs are also involved in planarian ERK activation and if so which cell types express them.

An outstanding question is whether the biochemical and possibly physical interactions that give rise to *Hofstenia*’s spreading ERK pattern are shared with other instances of ERK spreads or waves, phenomena in which ERK is sequentially activated in one cell, then its neighbors. In ERK waves in collectively migrating mammalian epithelial cells as well as the *Drosophila* tracheal placode, release of EGF family ligands from the cell membrane by protease-driven ectodomain shedding is required for waves (Ogura et al., 2018). Like other EGF family ligands, in mammalian cells NRG1 is released from the membrane via ectodomain shedding (Mei and Xiong, 2008). In light of this, one hypothesis as to how injury and *nrg-1* relate in *Hofstenia* is that injury triggers release of Nrg1 from muscle cell membranes via protease cleavage. This would increase extracellular Nrg-1 levels, and therefore Nrg-1-receptor binding and ERK activation. It is also possible, however, that Nrg-1 is not dynamically released from muscle cell membranes upon injury but instead is released homeostatically and already present in the extracellular space at the time of injury. Another open question is whether positive feedback plays a part in generating *Hofstenia’*s spreading ERK pattern. As other authors have proposed in the context of ERK waves, positive feedback could occur without requiring new gene expression e.g. if pERK positively regulated a protease responsible for Nrg-1 release, a relationship demonstrated for other EGF family ligands (Aoki et al., 2017; Deguchi et al., 2024; Díaz-Rodríguez et al., 2002; Fan et al., 2023; Hino et al., 2020; Murai et al., 2006; Umata et al., 2001).

Altogether, our work uncovers an important role for spreading ERK signaling, mediated by an EGF family ligand, during early stages of regeneration in an acoel. This work reveals similarities to mechanisms in vertebrates and provides advances beyond currently known mechanisms in systems with whole-body regeneration. These findings will serve as the basis of future studies of the mechanism of ERK regulation, such as how *nrg-1* signaling is triggered upon injury and whether positive feedback is needed to generate spreading, which will further enable comparative studies for understanding the evolution of regeneration.

## METHODS

### Animal maintenance

Adult animals were kept in artificial seawater (37 ppt, pH 7.9-8.0) at 21° C in plastic boxes within an incubator. Animals were fed freshly hatched brine shrimp (*Artemia*) twice per week, and water was changed and boxes cleaned twice per week. Juvenile animals were used for experiments and were raised in petri dishes as embryos, then in zebrafish tanks as juveniles, and fed rotifers (*Brachionus plicatilis*) twice per week.

### Small molecule inhibitor experiments

The inhibitors U0126 (Millipore Sigma 662005) and SU5402 (Sigma-Aldrich 0443) were resuspended from powder in DMSO to make concentrated stock solutions, which at the time of an experiment were further diluted in DMSO as necessary, then diluted in artificial seawater to a total DMSO concentration of 0.5%. For inhibitor experiments, animals were kept in 24 well plates with 1 ml liquid per well. For pre-amputation inhibitor treatments, intact animals were placed in a 24 well plate in a freshly prepared solution of the inhibitor in seawater. Animals were then amputated in a petri dish lid lined with filter paper and filled with additional fresh drug solution. After amputation, animals were placed back in the same well of the 24-well plate for post-amputation inhibitor treatment, in the same solutions used for pre-amputation treatment. For inhibitor washouts, 4 sequential washes with ample seawater were performed. Regeneration phenotypes of live animals were assessed using a Leica M80 stereomicroscope with mounted digital camera.

### RNAi interference (RNAi)

Double-stranded RNA (dsRNA) for injection (Srivastava et al., 2014) and soaking (Ramirez, 2020) were prepared using established protocols. The *C. elegans unc-22* gene, absent in *Hofstenia,* was used as a control gene. For assessing regeneration following *nrg-1* and *follistatin* RNAi, animals were injected with dsRNA for three days, then amputated at least two hours after injection on the last day. For the homeostatic *nrg-1* RNAi protocol used to assess ERK activation, animals were placed in soaking dsRNA in a 96 well plate. Animals were soaked for five days with injections performed on days 3, 4 and 5, then allowed to recover in seawater for 2 days. The second week, animals were soaked for an additional 5 days with injections again performed on the third, fourth and fifth day of that period. After the last day of injections, animals recovered in seawater overnight, then were amputated the following day and placed in soaking dsRNA until fixation. For the experiment in which muscle morphology was assessed, animals were soaked and injected for the first week as described above; the second week, animals were soaked for two days, then allowed to recover in seawater overnight and amputated the following day.

### Irradiation

Animals were subjected to 10000 rads, then amputated 7 days following irradiation. Previous work has shown that this leads to an absence of dividing cells by 7 days post-irradiation (Srivastava et al., 2014).

### Fixation and fluorescent in situ hybridization

Animals were fixed in 4% paraformaldehyde in phosphate-buffered saline with 0.1% Triton-X (PBST) or 4% paraformaldehyde in artificial seawater, for 1 hour at room temperature or overnight at 4 degrees. Following fixation, animals were washed 3 times with PBST. If only FISH was to be performed, animals were dehydrated in methanol. If IF or FISH with IF was to be performed, animals were stored in PBST at 4 degrees for up to 2 weeks. Digoxigenin and fluorescein FISH probes were prepared following an established protocol (Srivastava et al., 2014).

### Immunofluorescence staining

Animals were washed once with PBS for 10 minutes, then 4 times with PBST for 15 minutes each. Animals were blocked in 10% goat serum in PBST for 1 hour at room temperature, then incubated in primary antibody in block solution for two nights at 4° C. Animals were then washed once with PBS for 10 minutes, then 4 times with PBST for 15 minutes each, then blocked in 10% goat serum in PBST for 1 hour at room temperature. Animals were incubated in secondary antibody in block solution overnight at 4° C or for 3 hours at room temperature, then washed once with PBS for 10 minutes and at least 4 times with PBST for 15 minutes each. Animals were stained with Hoechst or DAPI 1:1000 at room temperature to label nuclei. Primary antibodies used are as follows: rabbit anti-pERK1/2 (Cell Signaling Technology 4370) 1:100, rabbit anti-phospho-histone H3 (Millipore 06-570) 1:100. Tropomyosin IF was performed using two custom primary antibodies following an established protocol (Hulett et al., 2020). For secondary antibodies, goat anti-rabbit Alexa 488, goat anti-rabbit Alexa 568, and goat anti-rabbit Alexa 647 (Abcam 150077, 175471, 150069) were used at a concentration of 1:1000.

### Image acquisition and quantification

All images of fixed animals were acquired using a Leica SP8 confocal and processed in ImageJ2 Version 2.14.0. For representative whole-fragment images, when necessary multiple tiles were stitched into a single image using the pairwise stitching plugin (Preibisch et al., 2009).

#### Image acquisition and quantification for pERK intensity profiles

For each fragment, an image encompassing the wound edge was acquired with the fragment’s anterior-posterior axis oriented vertically. Within an experiment, z-stack images were acquired with the same magnification, laser power, gain and z-slice interval. A maximum intensity projection was made of a fixed number of z-slices beginning with the first slice that lacked substantial epidermal staining. Next, images were processed using a custom pipeline that “de-wrinkles” the wound edge. First, a segmented line was drawn along the wound edge of the maximum projection image. If the exact position of the wound edge was not clear in the maximum projection, pERK and Hoechst signal in the original z-stack were examined, and the segmented line was drawn on that z-stack and then transferred to the projection image. The set of all coordinates along the segmented line was calculated. These coordinates were used by a script that takes every pixel (x,y) in the image and replaces its intensity value with the intensity value of the pixel at (x, y+*d*) where (x, *d*) is the coordinate of the wound edge line segment at x, effectively shifting each column of pixels upward by as many pixels as necessary to position the wound edge at y = 0. In the resulting transformed image, the wound edge is a straight line segment at the top of the image (y = 0). The resulting transformed images were manually inspected prior to analysis.

In each transformed image, a rectangle 200 µm wide was placed centered in x within the de-wrinkled area and with its upper edge at the upper edge of the image. The rectangle was extended as far away from the wound edge as possible given the boundaries of the transformed image, and without including any pERK staining in the brain or any artifactual staining of material in the gut. The plot profile command in Fiji was then used to obtain an average intensity profile along that rectangle’s y-axis. For a subset of fragments, a centered, 200 µm wide rectangle would have encompassed artifactual staining; for those fragments, the widest rectangle possible that did not encompass artifacts was used to plot a profile.

Further analysis of profiles was performed using Python. All profiles plotted and analyzed had a length above a cutoff specific to each experiment (350 µm for the pERK time course in Figure 1, 200 µm the SU5402 experiment in Figure 5 and 400 µm for the *nrg-1* RNAi experiment in Figure 5. To determine the location of each profile’s peak, profiles were smoothened using the pandas Series.rolling function with window size 10 pixels. The maximum value of the smoothened profile was called as the peak height, and the location of that value as the peak location. To determine the extent of above-background pERK signal, we used these smoothened profiles. We defined pERK extent as the minimum distance *d* from the wound site at which the absolute value of the profile’s slope fell below a specified threshold for two consecutive windows of slope calculation. We also specified that *d* must be greater than 60 µm for the wild-type time course (30 µm for the *nrg-1* RNAi experiment) because by visual inspection, all profiles’ extents were greater than these values in the respective experiments, and setting these conditions helped avoid incorrectly calling a local plateau within a profile as its extent.

The thresholds used were chosen for each experiment as the lowest slope for which most control profiles (or profiles at the latest time point) fall within that threshold at some distance; if a threshold slope too close to zero was chosen, many profiles’ slopes never fell below that threshold. With our chosen threshold for each experiment, a small number of fragments’ slopes never fell within the threshold range (2/57 for the time course in Figure 1, 3/50 in the SU5402 experiment in Figure 5), so a pERK extent could not be calculated. These fragments are included in the graphs showing average profiles as well as peak location and height, but excluded from graphs showing pERK extent.

#### Colocalization analysis

To determine the fraction of *tropomyosin^+^* muscle cells that are also pERK^+^, six fragments (three heads and three tails) per time point were imaged and analyzed using the Cell Counter plugin in Fiji. Images were acquired as z-stacks that encompassed the plane of body wall musculature (visible as bright, orthogonal *tropomyosin^+^* muscle fibers) and several planes interior to the body wall musculature, with 1 µm between slices. Images were rotated such that the wound edge was horizontal. All images in this analysis were acquired using the same imaging settings except for the total number of z-slices captured. The Gaussian blur command was applied to the pERK channel (radius 0.8). For *tropomyosin*, for each fragment, two rectangular areas 50 by 100 µm were analyzed: “area 1” beginning at the wound edge and extending 50 µm towards the anterior or posterior, and “area 2” spanning 50 to 100 µm away from the wound edge. For two fragments whose wound edge was curved or not orthogonal to the anterior-posterior axis, rather than one 50x100 µm rectangle per area, two 50x50 µm rectangles per area were analyzed. Within each area, *tropomyosin^+^* nuclei were marked. Then those nuclei were marked as either pERK^+^ or pERK^-^ by examining overlap of pERK and nuclear (Hoechst) signal, and pERK and *tropomyosin* signal, within multiple planes.

For *piwi-1*, pERK colocalization analysis, the same protocol was followed except that a 50 µm by 200 µm rectangular area was analyzed for each zone. For each rectangular area, counting was performed using nuclei visible within the z-plane that contained, by eye, the most *piwi-1^+^* cells.

### Quantitative PCR

Wound sites were isolated from amputated head and tail fragments, pooled and used for total RNA extraction with the Nucleospin RNA XS kit (Macherey-Nagel 740902.10). cDNA was prepared from extracted RNA using oligo dT-primed synthesis with the SuperScript III kit (Thermo Fisher 18080044). qPCR was performed using the SsoFast EvaGreen mix (Bio-Rad 172-5202). CT values were normalized to CT values for housekeeping gene EF1*α*. For each sample, a delta-delta CT (ddCT) value was obtained by subtracting the sample’s dCT value from the mean dCT value for 0 hpa samples under that treatment condition (DMSO or U0126). The ddCT values for 3 or 6 hpa samples were plotted and used for statistical testing. 3-4 biological replicates, each comprising 6-8 wound sites, were used for each treatment condition at each time point.

### Single-cell RNA-seq analysis

Expression of specific genes was visualized within existing intact and regenerating animal scRNA-seq datasets on UMAP projections and dot plots using Seurat v3 (Satija et al., 2015). The clustering used for each dataset was the clustering chosen through iterative testing of parameters, and validated with FISH, by (Hulett et al., 2023).

### Statistical analysis

Statistical tests were performed in Python using the SciPy stats module. Group means were compared using two-sample Welch’s t-tests, with or without assumption of equal variance as appropriate. Multiple testing correction was performed where stated using Benjamini-Hochberg false discovery control. Statistical significance was determined using a p-value cutoff of 0.05.

## ACKNOWLEDGMENTS

The authors would like to acknowledge Dr. Vikram Chandra for assistance with profile analysis and plotting, Carlos Rivera-López for providing feedback on the manuscript, and each member of the Srivastava lab for helpful discussion and feedback on this work. The authors also wish to acknowledge Dr. Douglas Richardson for advice regarding image quantification, as well as the Harvard Department of Statistics’ statistical consulting service.

---

**Supplemental figure 1.**
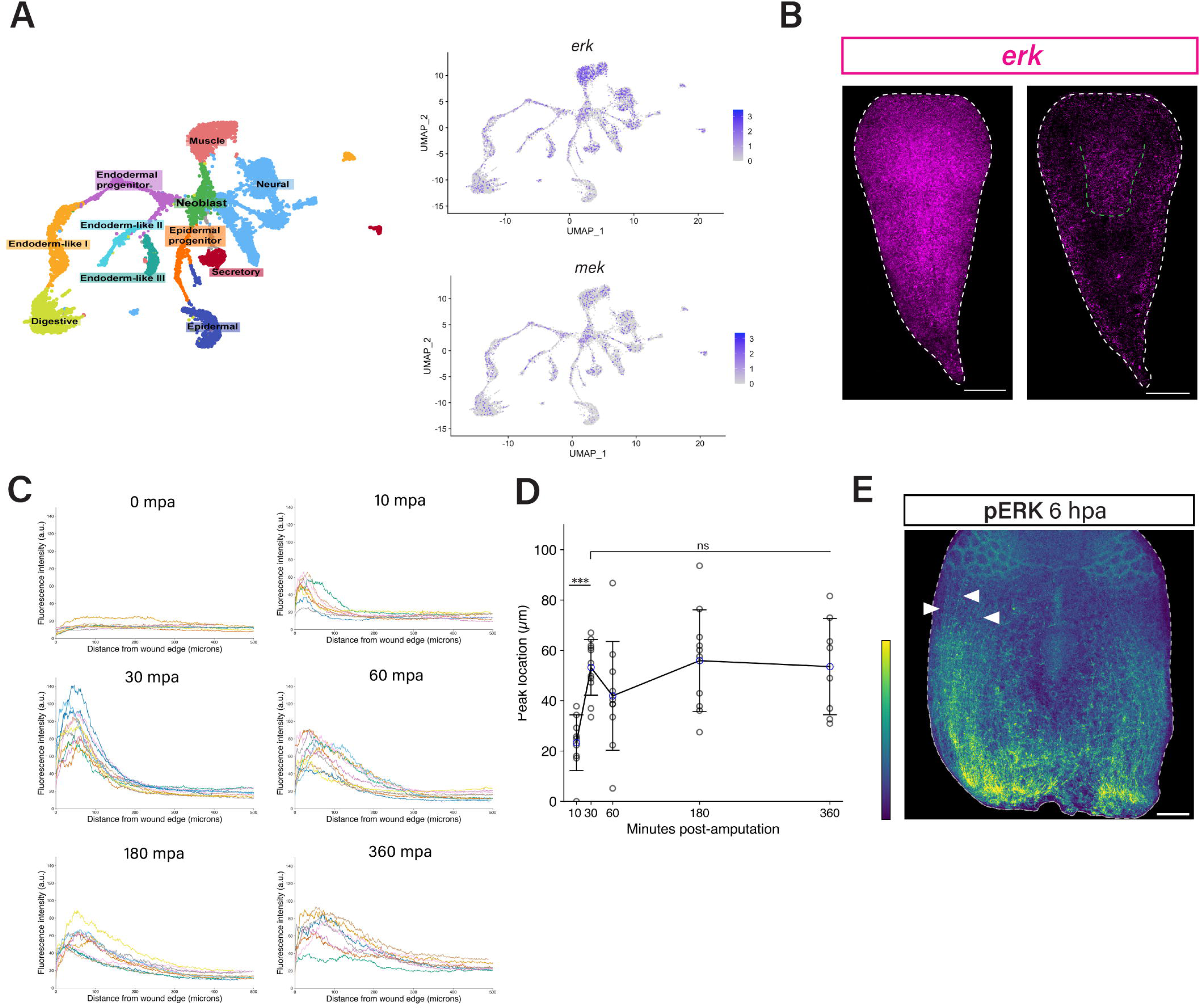
(A) Left, scRNA-seq data from (Hulett et al., 2023) showing cells in intact animals annotated with validated cluster identities. Right, projections of *erk* and *mek* mRNA expression in intact animals in these data. (B) *erk* expression in intact animals visualized using FISH. Left, maximum intensity projection of multiple optical slices including external and internal tissues; right, single optical slice within an internal plane. Green dotted line outlines the pharynx. (C) pERK profiles of individual fragments at each analyzed time point. Each individual profile is a different color. (D) Distance from the wound site of peak pERK intensity at 10, 30, 60, 180 and 360 mpa. Blue circles show means; error bars show standard deviation. *** p < .001, ns, not significant, two-sided Welch’s t-test. Adjusted p-values: 10 vs 30 mpa 4.29e-5, 30 vs 60 mpa 0.282, 30 vs 180 mpa 0.945, 30 vs 360 mpa 0.964. (E) 6 hpa head fragment with pERK^+^ muscle fibers distal to the wound site (arrowheads). Intensity scale in image is increased relative to scale used in Figure 1B. Scale bars, 100 µm.

**Supplemental figure 2.**
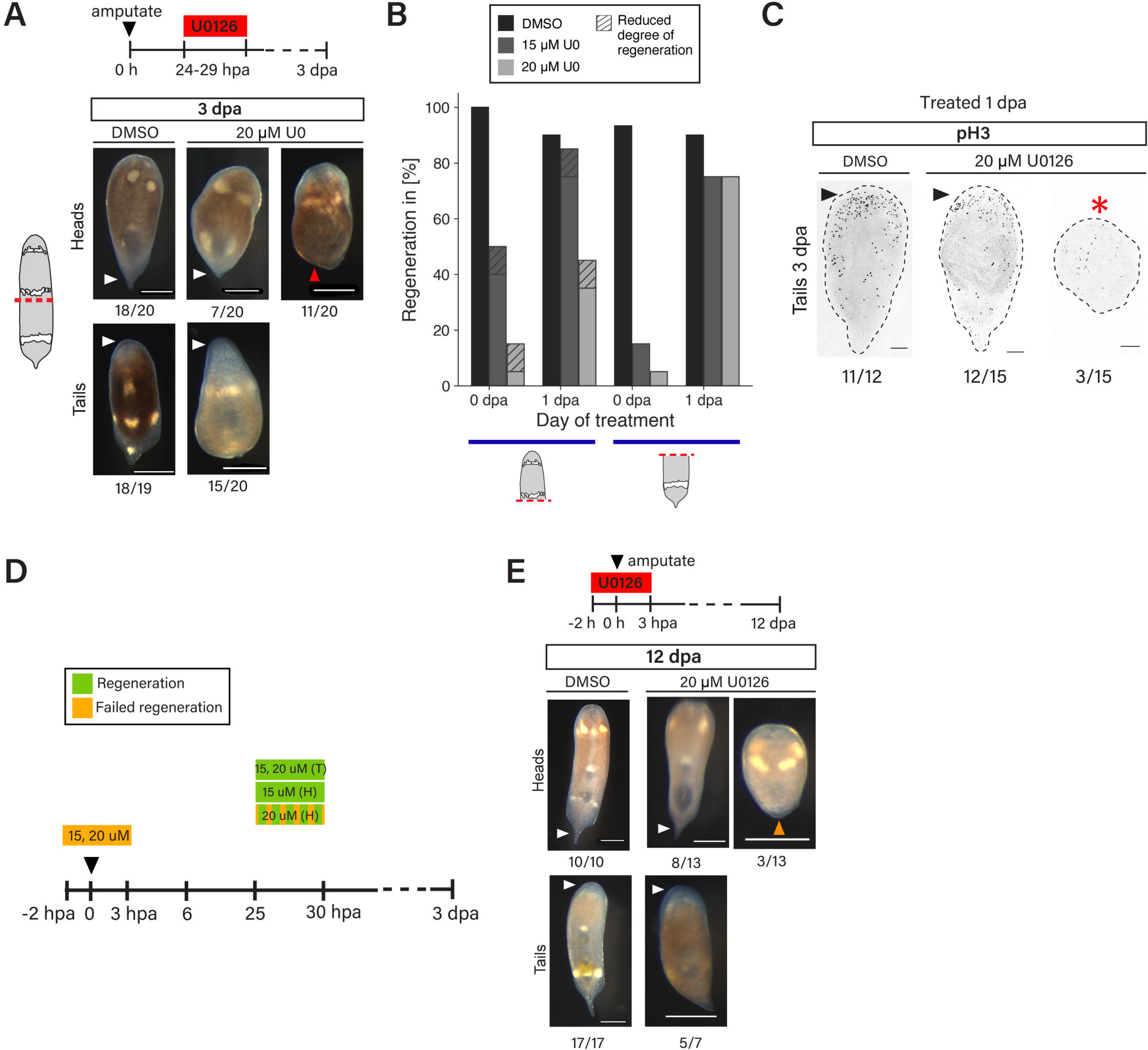
(A) Top, schematic showing timing of U0126 treatment for animals treated at 1 dpa. Bottom, regeneration outcomes for animals treated with DMSO or 20 µM U0126 at 1 dpa. Control (DMSO-treated) fragments regenerated a head or a tail as appropriate (white arrowheads). Among 20 µM U0126-treated head fragments, some regenerated a tail (white arrowhead) but more than half did not (red arrowhead). A majority of 20 µM U0126-treated tail fragments regenerated a head (white arrowhead). (B) Percentages of phenotypes observed in a single experiment in which animals were treated with U0126 at either 0 or 1 dpa (Figure S2A, Figure 2C). A subset of the 0 dpa-treated animals included in the bar plot in Figure 2D are also included in the counts represented in this plot. (C) pH3 IF in 3 dpa tail fragments following treatment with DMSO or U0126 at 1 dpa. Arrowheads, anterior accumulation of pH3^+^ cells. *, scattered pH3^+^ cells without clear anterior accumulation. (D) Schematic summarizing regeneration outcomes following U0126 treatment during different time windows. See also phenotype scoring data in Table S1. (E) Regeneration outcomes at 12 dpa. At 12 dpa, 8/13 U0126-treated head fragments had regenerated a normal tail (white arrowhead) and 3/13 showed visible but reduced new tissue formation (orange arrowhead). Among U0126-treated tail fragments, 5/7 had formed a visible head blastema (white arrowhead) and 2/7 showed no head blastema. Animals shown were also part of phenotype scoring at 3 dpa shown in (B) and Figure 2C-D. Scale bars, 100 µm (C), 300 µm (A, E).

**Supplemental figure 3.**
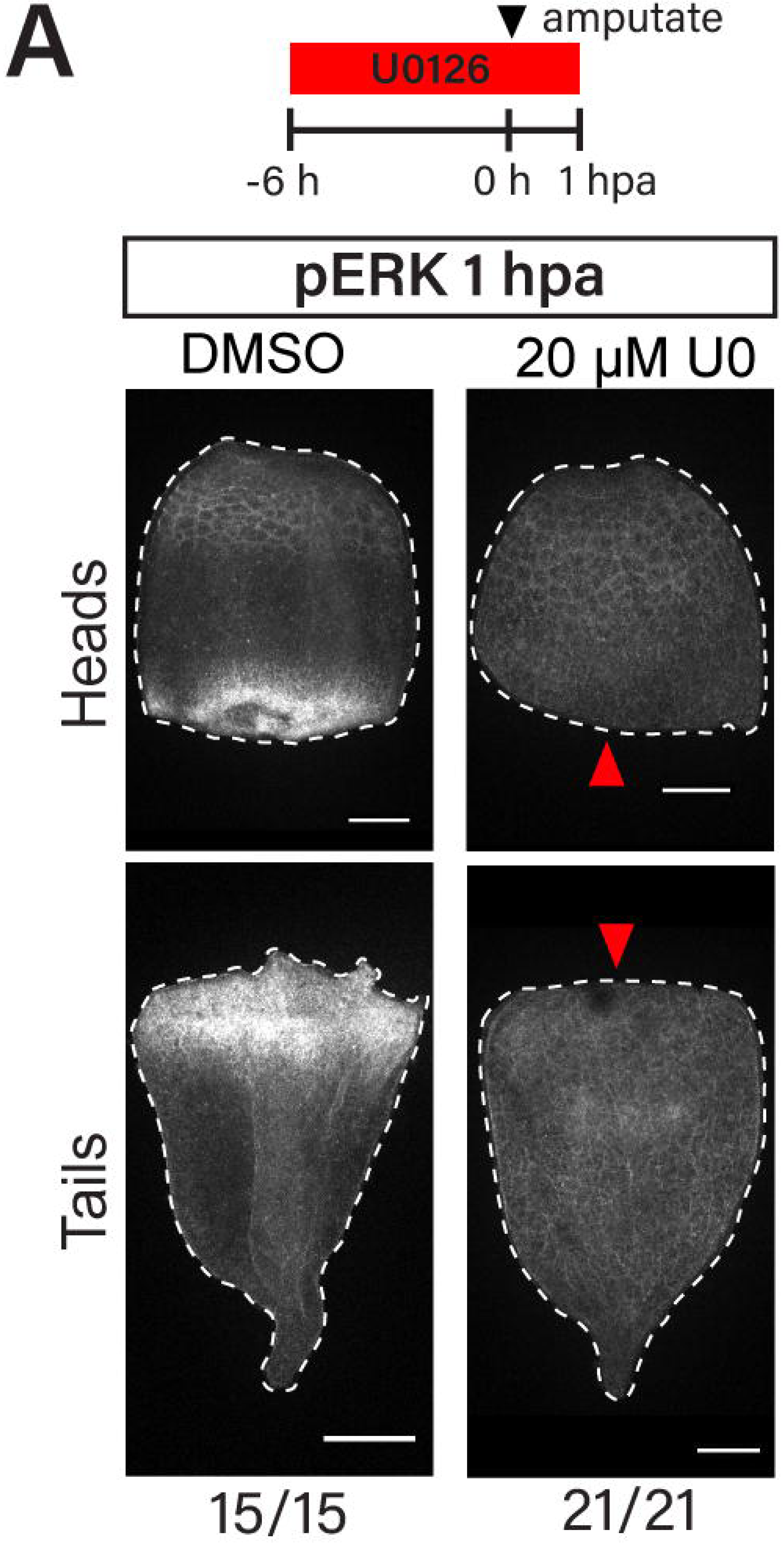
(A) Top, schematic showing timing of U0126 treatment. Bottom, pERK IF at 1 hpa in animals treated with 0.5% DMSO or 20 μM U0126. 20 μM U0126 treatment robustly inhibits ERK phosphorylation at wound sites at 1 hpa. Scale bars, 100 µm.

**Supplemental figure 4.**
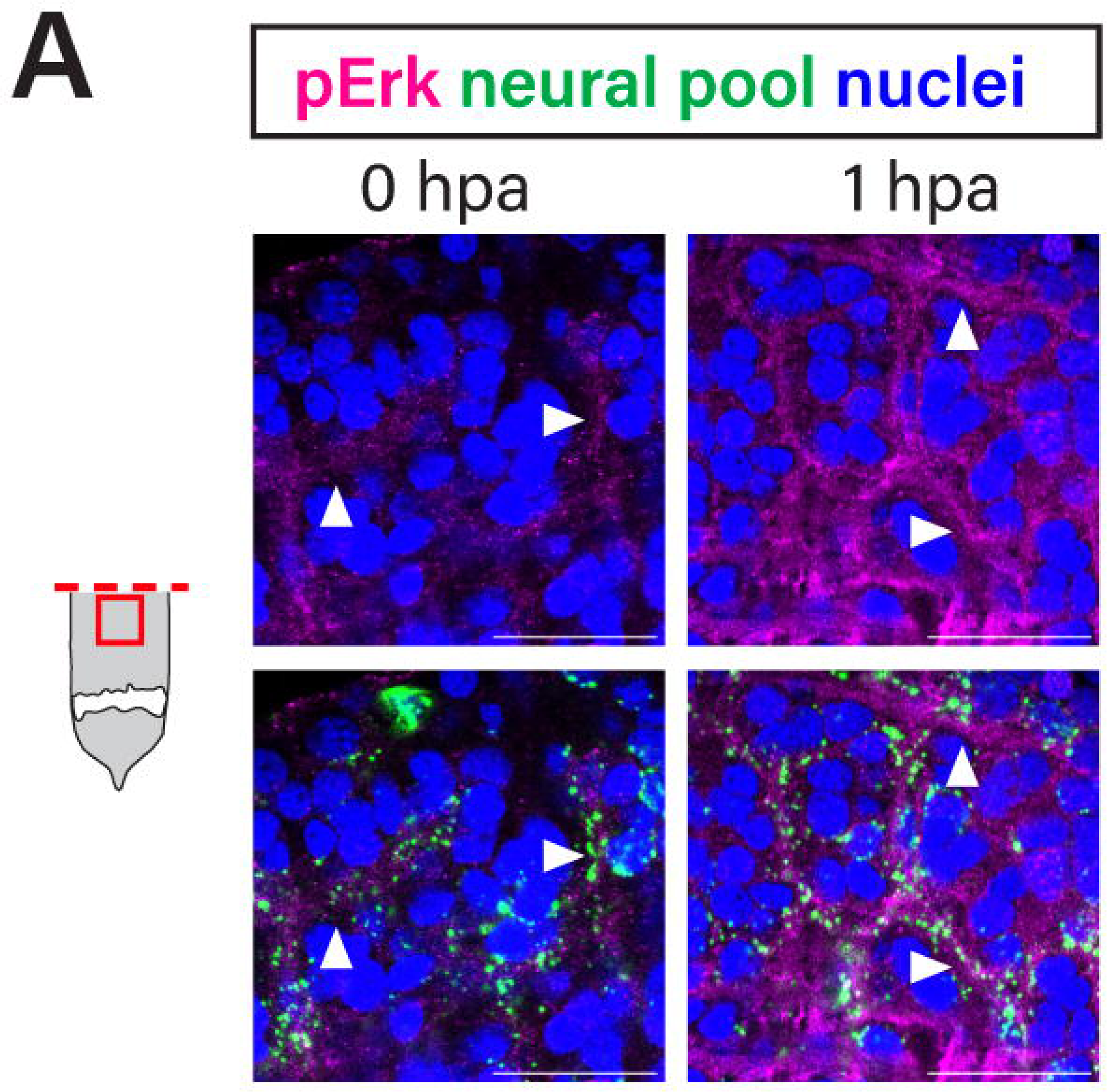
(A) FISH for neural marker pool combined with pERK IF. Images shown are of wound-adjacent region of a tail fragment. As in head fragments, pERK signal within neurons in the wound-adjacent region appears to increase in intensity between 0 and 1 hpa. Scale bars, 20 µm.

**Supplemental figure 5.**
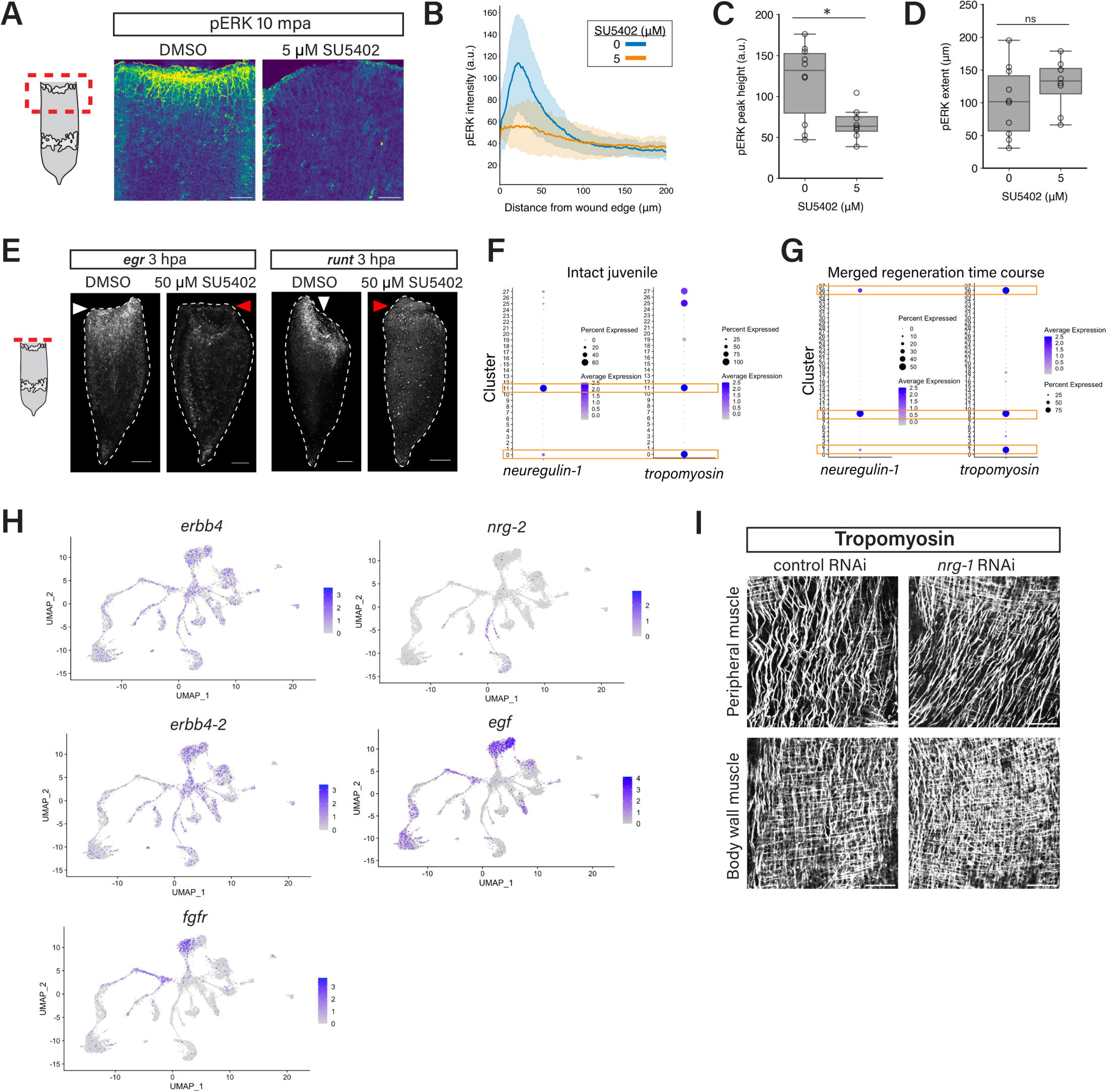
(A) pERK staining at 10 mpa in representative tail fragments treated with 0 (DMSO) or 5 μM SU5402. (B-D) Average pERK profiles (B), profile peak height (C) and pERK extent (D) in control and 5 μM SU5402-treated fragments. pERK peak height is significantly different between groups (p-value = .0104) but pERK extent is not (p-value = 0.523). (E) *egr* and *runt* expression at 3 hpa in tail fragments treated with DMSO or 50 µM SU5402. 3/3 SU5402-treated tails showed sharply reduced *egr* expression as compared to controls (red arrowheads; 9/9 control tails with normal *egr* expression, white arrowheads). 6/6 SU5402-treated tails showed reduced *runt* expression as compared to controls (red arrowheads; 5/5 control tails with normal *runt* expression, white arrowheads). (F, G) scRNA-seq dot plots (data from (Hulett et al., 2023)) showing *nrg-1* and *tropomyosin* expression in cell clusters of intact animals (F) and animals in a regeneration time course (G) including 0, 6, 24 hpa, 3 dpa and later time points. In (G), clusters with high *nrg-1* expression (boxes) also have high *tropomyosin* expression. (H) Projections showing expression in intact animals of putative ERBB homologs *erbb4* and *erbb4-2,* a putative FGF receptor (*fgfr*), and putative EGF family ligands *nrg-2* and *egf.* (I) Body wall muscle and peripheral muscle of homeostatic *nrg-1* RNAi animals showed no gross morphological defects (n = 9/12, control n = 10/12). Shading in (B) and error bars in (C, D) show standard deviation. * p < 0.05, ns, not significant, two-sided Welch’s t-test. Scale bars, 50 μm (A, I), 100 μm (E).

## SUPPLEMENTAL TABLES

Table S1. Live phenotype scoring data for all regeneration assays

